# Software Evaluation for *de novo* Detection of Transposons

**DOI:** 10.1101/2021.02.08.430290

**Authors:** Matias Rodriguez, Wojciech Makałowski

## Abstract

Transposable elements (TEs) are major genomic components in most eukaryotic genomes and play an important role in genome evolution. However, despite their relevance the identification of TEs is not an easy task and a number of tools were developed to tackle this problem. To better understand how they perform, we tested several widely used tools for *de novo* TE detection and compared their performance on both simulated data and well curated genomic sequences. As expected, tools that build TE-models performed better than k-mer counting ones, with RepeatModeler beating competitors in most datasets. However, there is a tendency for most tools to identify TE-regions in a fragmented manner and it is also frequent that small TEs or fragmented TEs are not detected. Consequently, the identification of TEs is still a challenging endeavor and it requires a significant manual curation by an experienced expert. The results will be helpful for identifying common issues associated with TE-annotation and for evaluating how comparable are the results obtained with different tools.

## Introduction

The vast majority of eukaryotic genomes contain a high number of repetitive DNA sequences. These sequences can be broadly classified as tandem repeats or interspersed repeats. Tandem repeats are short sequences with a length up to a few dozen bases that lie adjacent one to another and are approximate copies of the same pattern of nucleotides. Similarly, interspersed repeats are homologous DNA sequences that can be found in multiple copies scattered throughout a genome and their lengths can vary immensely from a hundred nucleotides up to more than twenty-thousand nucleotides. Most of these interspersed repetitive sequences found in genomes originated from the proliferation of transposable elements.

Transposable elements (TEs) are mobile genetic sequences possibly related to viral components that have evolved the ability to increase their abundance in a genome by making copies of themselves. The fraction of TEs in a genome can vary widely and can represent more than 80% of plant genomes (Schnable et al. 2009). To put into perspective how common they are, if we consider a well-studied case, such as the human genome, the annotated protein-coding genes represent only a very small fraction of approximately 5% of all the sequences, meanwhile TEs can make up to about 68% of the sequences (de Koning et al. 2011).

Genomes and TEs have coevolved similarly to a host-parasite relationship and this led the genomes to develop multiple mechanisms to suppress TE activity as they can compromise the integrity of the genome and can cause deleterious mutations. Consequently, there is a constant evolutionary arms race between transposon activity and the host genome trying to suppress their proliferation (Jurka et al. 2007). Despite the parasitic nature of TEs, they play a fundamental role in genome evolution, contributing to plasticity, shaping, and altering the architecture of the genome. TEs contribute to gene regulatory networks as their activity can disrupt regulatory sequences modifying gene expression by altering chromatin structure, behaving as enhancers or promoters, or, when transcribed as part of a larger transcript, creating new transcript isoforms altering splicing and mRNA stability (Makalowski 2000). There are multiple examples of TEs that have been domesticated and proteins derived from them which were co-opted, such as the RAG1 gene from the somatic V(D)J recombination in humans and the retrotransposons that maintain the telomeres in *Drosophila*. RNA-mediated retrotransposition of transcribed genes is also a source for gene duplications that can lead to novel traits (Kubiak and Makalowska 2017).

Historically, TEs were considered useless selfish sequences and their influence on genes and genomes was often dismissed (Ohno 1973). It was not until the last two decades that they started to be considered as major components of genomes and important players of genome evolution, but due to the difficulties posed by their repetitive nature their annotation and role in genetic studies still continues to be neglected (Biemont 2010).

The correct identification of TEs is an important step in any genome project since their repetitive nature can create difficulties during *de novo* genome assemblies, breaking the continuity of contigs as a result of the same reads mapping to multiple loci (Ricker et al. 2012). They can also hinder annotation by creating conflicts with gene prediction programs if they can be found inside a host gene, carry part of a host gene when replicating, become pseudogenes, or contain spurious ORFs.

There are multiple tools for TE-detection but there are no clearly defined pipelines or software tools that could be considered as standard, as there are no clear metrics to compare the results obtained from each software (Hoen et al. 2015). Most tools also rely on a high copy number of elements for correct identification and are usually tested in organisms that have large genomes and a high abundance of TEs.

The identification of TEs can be a real daunting and a time-consuming endeavor for the amount of data that needs to be processed and compared and the challenges inherent to their complex nature. TEs are extremely diverse, they comprise multiple classes of elements that can vary immensely in sequence, length, structure, and distribution (Wicker et al. 2007). Some TEs families found in eukaryotic genomes can be very old with a majority of inactive copies due to accumulation of mutations or fragmentation during the insertion process. This means that remains of antique copies from a family can be very divergent from active TEs, making the detection of the remnants of decayed copies or the definition of consensus sequences a real challenge that is hindered by the great variability of TEs within the same family. The proliferation of TEs can also result in the generation of nested TEs and some families show a clear preference for jumping into other TEs that act as hotspots for insertion (Gao et al. 2012), making the detection and correct annotation of them even more difficult.

There are well curated TE databases, such as RepBase (Bao et al. 2015) or Dfam (Hubley et al. 2016), with libraries of consensus sequences. A homology-based approach relies on the TE sequences from these libraries which are then mapped against the studied genome. To identify new TEs a *de novo* approach is used and there are abundant software alternatives which rely on different strategies ranging from structural information, periodicity, k-mer counting, or repetitiveness, among others (Makalowski et al. 2019). When a new species is sequenced, a strategy which uses only information from curated databases is not enough and it is necessary to use a *de novo* strategy to identify novel families and species-specific TEs.

In this work we compare TE detection software which are widely used by researchers and we assess their performances on genomes with well curated TE annotations. We ran a number of *de novo* TE detection software packages on simulated sequences and genome sequences and then compared and evaluated their performance in detecting a wide variety of divergent TE families. A particular scenario that we tried to consider is the detection of transposons in smaller genomes of around a hundred million bases. In all cases the software for identification of TE rely on the presence of a large number of elements of the same family and that is usually not common in smaller genomes where a lower number of copies of TEs is expected.

## Methods

### Datasets

Genomic data with annotated TEs were downloaded from the UCSC Genome Browser database (Haeussler et al. 2019). The TE annotation provided by UCSC Genomes was obtained from mapping TEs from the RepBase database (Bao et al. 2015) against each genome using RepeatMasker (Smit et al. 2013-2015). The sequence data sets we used varied from 46.7 Mb for the human chromosome 21 and 137.5 Mb for the fruit fly genome (see Table 1).

**Table 1.** Datasets used for testing the TE *de novo* detection tools.

| Dataset | Genome version | Sequences | Length | TE-fraction |
| --- | --- | --- | --- | --- |
| Human genome | GRCh38/hg38 p12 | Chromosome 21 | 46.7 Mb | 36.74% |
| Zebrafish | GRCz11/danRe 11 | Chromosome 1 | 59.6 Mb | 46.58% |
| Fruit fly | BDGP Release 6 | Whole genome | 137.5 Mb | 17.55% |
| Simulated data | n/a | Simulation | 100.1 Mb | 60.00% |

### Simulated data

We used a Python script to simulate an ideal scenario where the composition, coordinates, and divergence of all the TEs are already known. This script takes input from a configuration file for GC content, TE sequences, number of copies, expected divergence (mutations and indels), percent of fragments, and nesting. The script starts by simulating a random sequence of a predefined length and a GC content that constitutes the base sequence where TEs are going to be inserted. Then it obtains the name of each TE and the number of copies from the configuration file and random positions are chosen and assigned to each TE. In the next step, TE sequences are loaded from a library and the information about divergence and fragmentation is taken into account to generate random mutations and fragments that are inserted into the base sequences. The last step takes all of this information to generate a fasta file with the whole new sequence with TEs and a GFF file with all the coordinates and relevant information. For the simulated dataset we used a base sequence of 40 Mb and inserted 60 Mb of TE-sequences from 20 different families downloaded from the Dfam database (Hubley et al. 2016). Although the divergence threshold for individual copies inserted into simulated genome was set to 30%, the majority of the sequences were between 90% and 95% identical to the cognate TE consensus sequences (see Figure 1). Detail information on inserted sequences is provided in Table S1.

**Figure 1.**
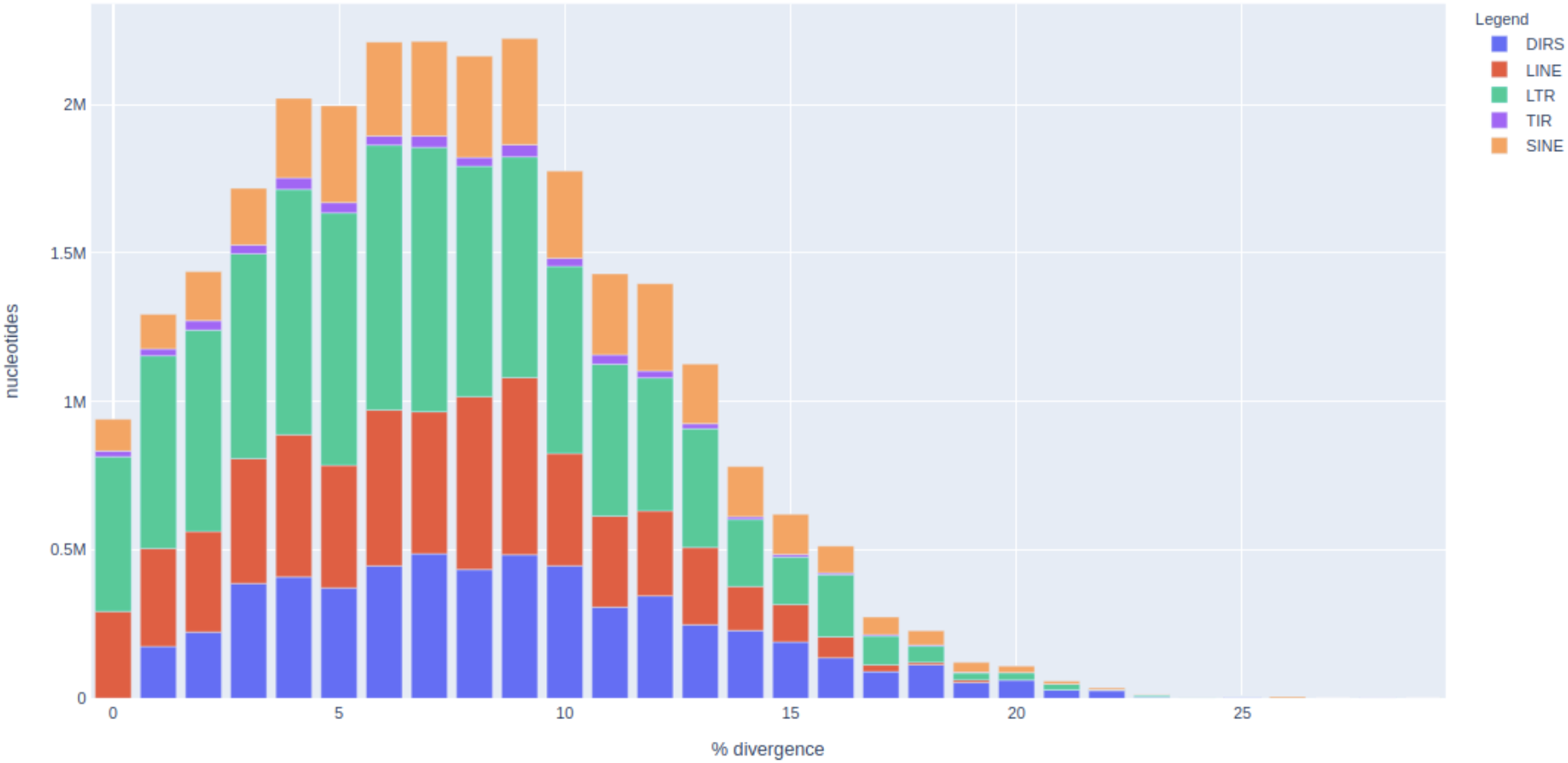
Landscape plot of simulated TE insertions.

### Software

In this work we compared strategies for *de novo* detection of TEs using k-mer based tools and programs that construct TE-models. We tested three k-mer counting tools: Red, P-Clouds, and phRaider. Due to the nature of the algorithms employed by k-mer counting software, these tools are extremely fast and usually don’t require much computational power. Nevertheless, they usually require a big amount of RAM to store data structures, so they may not scale up well with large genomes. Red identifies candidate repetitive regions giving them a score, then processes these results using signal processing and the second derivative. These filtered data are used to train a Hidden Markov Model that scans the genome for candidate TEs. As described by the author, it is a novel repeat discovery system that trains itself automatically on an input genome (Girgis 2015). P-Clouds counts oligonucleotides, then arranges them into clusters of #probability clouds” that are related oligonucleotides that occur as a group more often than expected by chance. Then it annotates the genome by finding stretches with a high density of oligos present in these “probability clouds” (Gu et al. 2008). The premise of phRAIDER is to use spaced seeds to specify match patterns, i.e., to permit the search of substrings allowing mismatches in certain positions. Then it scans the genome searching for highly frequent seeds and how they overlap (Schaeffer et al. 2016). The biggest limitation of these tools is the fact that they don’t make any attempt to classify found repeats. Moreover, there is no information provided on relation between the detected individual elements and no consensus sequences are computed.

We also compared three model-builders: RepeatScout, REPET (TEdenovo pipeline), and RepeatModeler. RepeatScout uses high frequency seeds and extends each seed to a progressively longer consensus sequence, following the dynamically inferred alignments between the consensus sequence and its occurrences in the genome. The alignment score encourages boundaries shared by some but not necessarily all alignments; it uses a standard SW-algorithm to extend until n-iterations fail to improve the score (Price et al. 2005). REPET is a package consisting of two pipelines, one for detection of TEs: TEdenovo, and another for their annotation: TEannot (Quesneville et al. 2005; Flutre et al. 2011). Both of these pipelines are fully configurable and each step can be parametrized. The TEdenovo pipeline by default starts self-comparison of the input genome with BLASTER, a modified version of BLAST. Then it clusters the high scoring pairs using three tools: RECON, GROUPER, and PILER, grouping closely related TE sequences. Finally, it performs a multiple alignment using MAFFT or MAP with the aim of having a consensus sequence for each TE family. Here, we are interested in the ability of the software to detect TEs, so we used only the TEdenovo pipeline. Finally, RepeatModeler is a pipeline that uses as an input the outputs of three other software, namely RECON, RepeatScout, and Tandem Repeats Finder. Additionally, it uses LTRHarvest or LTRretriever for LTR-TE detection (Flynn et al. 2020). It should be noted that RepeatModeler contains an optional module (option -LTRStruct) that enables clustering redundant LTR models. However, since we used only default parameters, this step was skipped in our analyses. RepeatScout, RepeatModeler, and REPET all give as a final result a fasta file with a consensus sequence for each type of TEs they could identify. Afterwords, we need to map back these consensus sequences to the genome to identify individual copies of TEs and get their coordinates. For this step we used the popular tool RepeatMasker, version 4.0.9 (Smit et al. 2013-2015), using the three tools’ outputs as libraries to screen the genomes for TEs.

Another tool used for detecting simple repeats is Tandem Repeat Finder (TRF) (Benson 1999), a software which models tandem repeats using a probabilistic model. We used it to filter out simple repeats obtained by the k-mer counting tools and also to assess the ability of model-builders to cope with simple sequence repeats. TRF was run with default parameters on all the sequences analyzed and the results obtained were merged when an adjacent or overlapping annotation was reported. These results were converted into a GFF file for easing further analysis.

### Pipeline

All the software were tested with default parameters as we intended to compare the average performance of each tool without tuning their optional parameters (see Figure 2). K-mer counting methods are expected to find all the high frequency k-mer, including simple repeats and interspersed repeats. For k-mer counting software after getting the results we ran Tandem Repeat Finder (TRF) with default parameters to filter out tandem repeats. The model-builders usually include some steps for filtering simple repeats and frequently make use of TRF, so this step was not replicated with these software.

**Figure 2.**
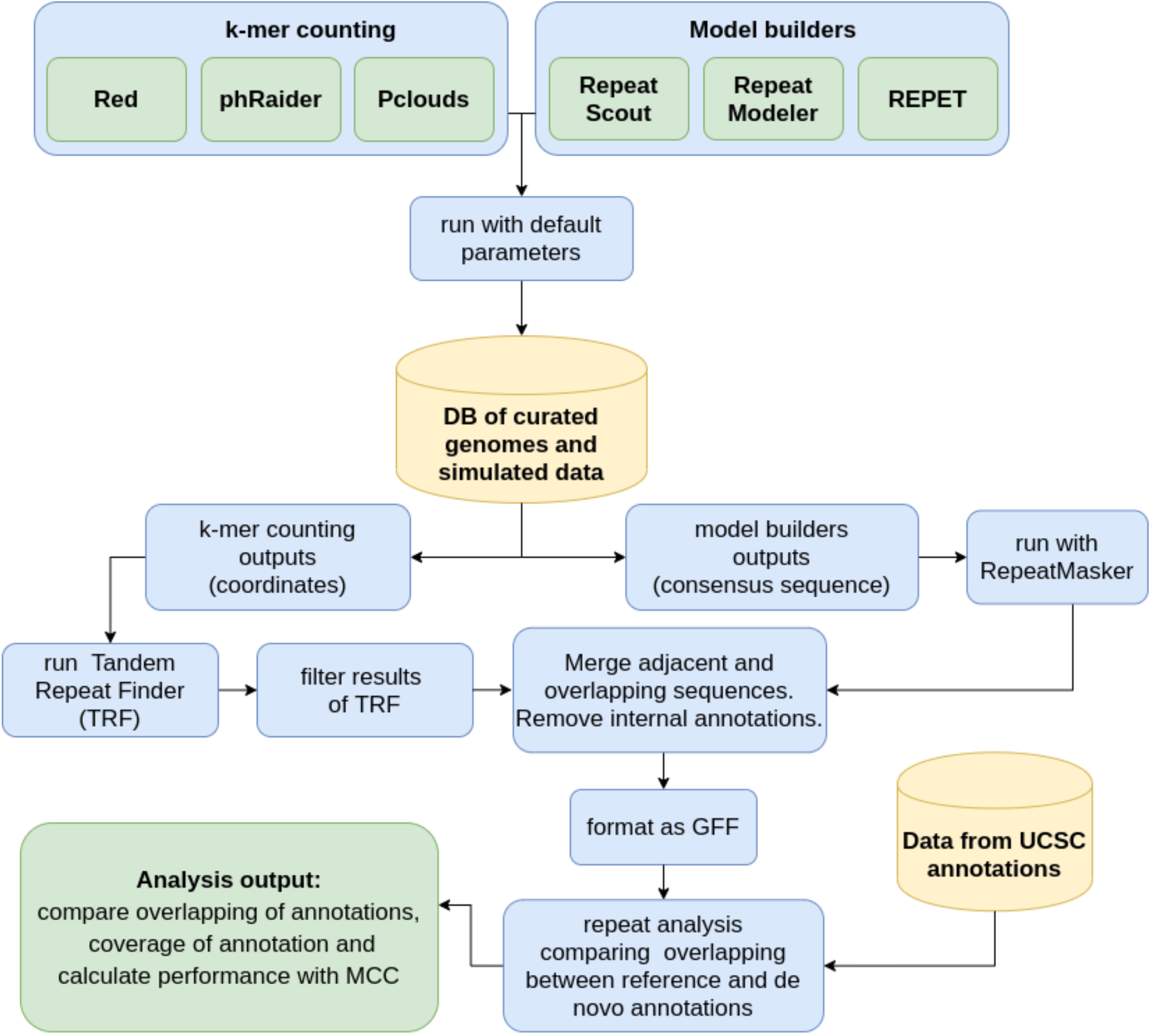
Pipeline used for testing and comparing the performance of *de novo* detection tools.

The type of results produced by each set of tools is also quite different due to the different strategies used. K-mer counting tools return the coordinates in the genome with the regions where high frequency k-mers were found, meanwhile model-building software returns the consensus sequences of the TE candidates found as a fasta file. So for the latter set of tools we mapped back the consensus sequences against the original sequences using RepeatMasker, running it with default parameters. The results obtained from all the different tools were transformed to GFF format for further processing.

Then GFF files were sorted by coordinates and immediately adjacent, overlapped, or internal coordinates were merged as one, as the main idea is only the identification of transposon sequences. This step is necessary particularly in k-mer counting software which have the tendency to annotate many overlapping and fragmented repeats. For all the datasets tested we have as a reference the annotations downloaded from the UCSC Genomes database.

### TE-models’ comparison

TE-models generated by model-building software on simulated data were compared to original TE-families using blastn (version 2.10.1+) with default parameters.

### Analysis

With the results of TE detection obtained from each software we created GFF files that were then compared against the original files from UCSC database with RepeatMasker mapping results. A custom Python script was used to obtain the overlapping regions of two GFF files and where there are differences in the annotation, it allowed us to compare the coordinates of the reference and the ones obtained by the TE *de novo* software. This way of evaluating the data is useful in order to create a confusion matrix that can be used as input for a binary classifier test that allows us to compare the performance of different software against a reference. When the reference annotation and the new annotation agree on the coordinates, these bases are counted as true positives, or if nothing is annotated in both, these bases count as true negatives (Figure 3). If the new annotation has bases not covered by the reference annotation, we consider them as false positives, and similarly if annotations in the reference are missing in the new annotation, these are counted as false negatives (see Figure 3).

**Figure 3.**
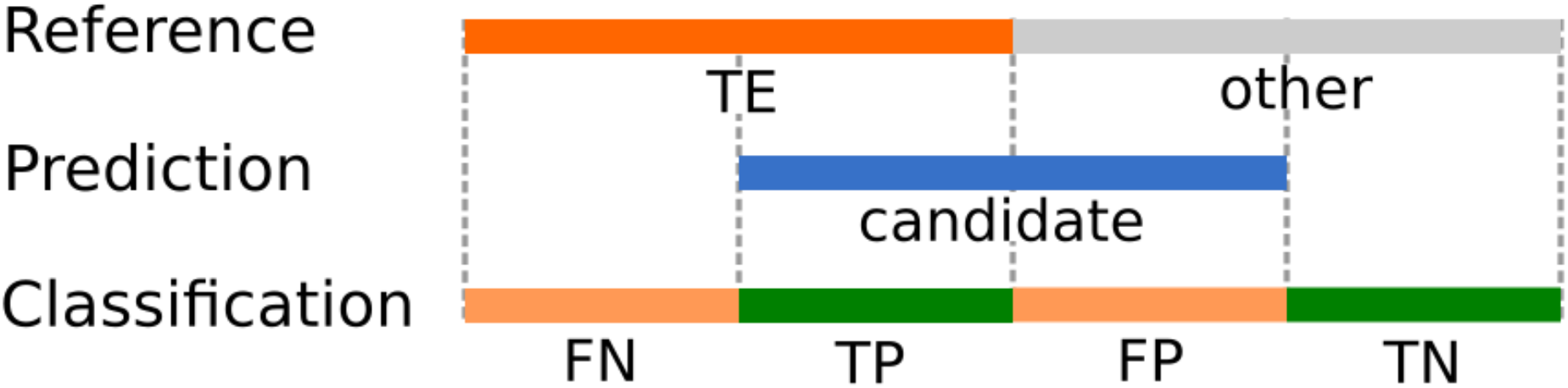
Classification of the results obtained after comparing the reference annotation and the predicted TEs. We have TEs in the reference genome annotation and TE candidates as a prediction. Then comparing both they can be classified as false negatives (FN), true positives (TP), false positives (FP), or true negatives (TN).

With this kind of data we have a binary classification problem, where each category can be classified using a confusion matrix. There are multiple tests to evaluate and compare the results obtained by a binary classifier which make use of a confusion matrix data and one of the most commonly used methods is the Matthews Correlation Coefficient (MCC). The MCC has the advantage that it uses all four quadrants of a confusion matrix considering the proportions of each class and requires that in both classes negative and positive elements are correctly classified, performing well even when using imbalanced data and when one class is underrepresented (Boughorbel et al. 2017). The MCC evaluates the results obtained from a prediction, as in this case the TE *de novo* software TE candidates, against the known annotated data. The values of MCC range from −1 to +1, where a value of −1 is obtained when all the predictions are wrong, 0 when results are not better than random guessing, and 1 where all predictions are correct. In this work we used the MCC as a measure of the performance obtained from the different software tools. Additionally, we developed several R scripts for plotting GFF coordinates which visually compare the annotations obtained from each tool.

## Results

As mentioned above, the two groups of programs provide different types of results. While k-mer counting software provided a list of regions that are occupied by repetitive sequences, model-building software analyzed here returned sets of repeats’ models. These models can be next used to scan a genome and annotate individual repeats, including TEs. For this step, we used a popular program, namely RepeatMasker (see Methods section).

### Model building

Three different programs were used to create TE-models for both real genomic data and simulated sequences. The results of the latter are the most informative as we knew the exact number of expected TE-families. Interestingly, all three programs generated more TE-models than we used for the simulation (see Table 2).

**Table 2.** Number of TE models generated by each software from a simulated sequence containing TEs of twenty different families (see Table S1 for detailed information). Please note that REPET failed to generate an Alu model.

| Software | Number of models generated |
| --- | --- |
| RepeatScout | 30 |
| RepeatModeler | 80 |
| REPET | 82 |
| Number of TE families inserted in the simulated data | 20 |

RepeatScout generated the smallest number of TE-models (30) and only in few cases more than two models for a given TE-family: three for L1 and four for Polinton. However, it has a tendency to create homo-dimeric elements, for instance Copia, DIRS, HERVL (see Table S1). On the other extremum lies REPET, which created the highest number of models, although it failed to report an Alu model. This is a bit surprising since there were 730 Alu insertions of a single sub-family (Alu Y). REPET not only generated the highest number of models but some of them were dimeric and hybrid. The latter were caused by a few nested repeats, for instance a Jockey nested in a Ngaro or a Tc1-Mariner nested in a HERVL. In general, longer elements tend to give rise to several models by both RepeatModeler and REPET. For instance, 5.5 kb long L1 element is a source of six models in RepeatModeler analysis and nine models in the case of REPET. Polinton, which is 18.5 kb, resulted in eight models in both RepeatModeler and REPET and four models in the case of RepeatScout (see Table S1). Interestingly, some of these models overlap each other, suggesting that they could be merged during manual curation.

In our simulation, we “mutated” individual TE-copies up to thirty percent divergence from the reference sequence and many of the individual copies were truncated at the 5’ end. In general, a consensus sequence recovery at the nucleotide level was very good, with the average sequence identity of models to their respective reference sequences at 97.3% (stdev = 3.76). However, many of TEs were broken by a given software into several models. Probably the best example is Polinton, based on which RepeatModeler and REPET created nine models each and none of these covered the whole Polinton sequence that was used in the simulation (see Figure 4). The shortest transposon inserted into the simulated data was the 311 nucleotide long AluY element. The individual sequences were “mutated” to average 13% divergence from the reference sequence and 67 of them were truncated at their 5’ end up to 30% of the sequence length. RepeatScout performed the best, returning a 308 nt long model with the sequence identical to the reference and just 3 nt missing from the 5’ end. Surprisingly, REPET didn’t report any models based on these sequences. Finally, RepeatModeler created three different models: one almost ideal with just a 5’ terminal guanosine missing and two others, a bit shorter but with extra eight and thirty-five nucleotides added to their 5’ end (see Table S1).

**Figure 4.**
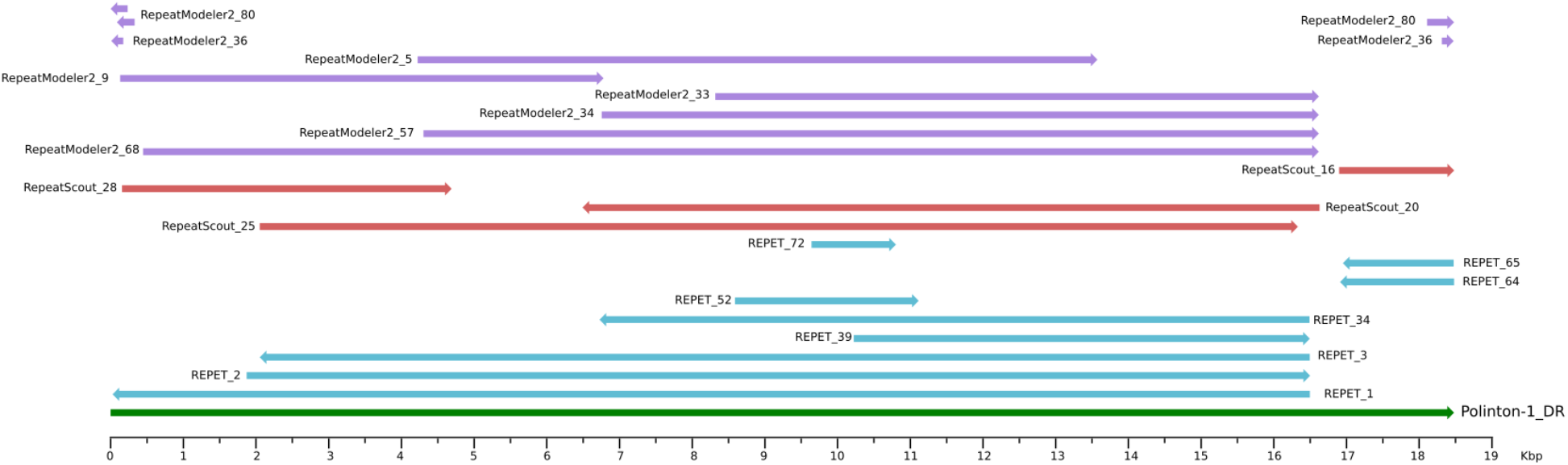
Different models created for Polinton-1_DR transposon aligned against Dfam model DF0002823.2.

When analyzing the simulated data, one trend became clear, namely that on occasions multiple models are generated for the same TE and there are common patterns observed for each software. A characteristic of RepeatModeler is that it tends to generate redundant models, with up to six to eight models for the longest TEs. This is also a common behavior observed with REPET, where many fragments were generated. Another interesting observation that only occurs with REPET is that in some models part of a nested TE was included into a model resulting in chimeric models. With RepeatScout there is much less redundancy with the number of models, but again something unique happens and some of the reported models are total or partial duplicates of the original TE.

An example of different models generated for a DIRS TE of 5.6 kb which were particularly difficult to resolve is shown in Figure 5. This particular TE RepeatModeler generated six different models of different lengths. These models are on average fifteen percent diverged at the sequence level. There were two models calculated by RepeatScout, one almost identical to the reference and another one almost twice the length of the original and consisting of a duplication leading to erroneous homodimer. Interestingly, the two copies of this homodimer are complementary to each other as they lie on opposite strands compared to the reference sequence. REPET reported four models in total. Two are relatively short, encompassing about one-third of the reference sequence and partially overlapping in a head-to-tail orientation.

**Figure 5.**
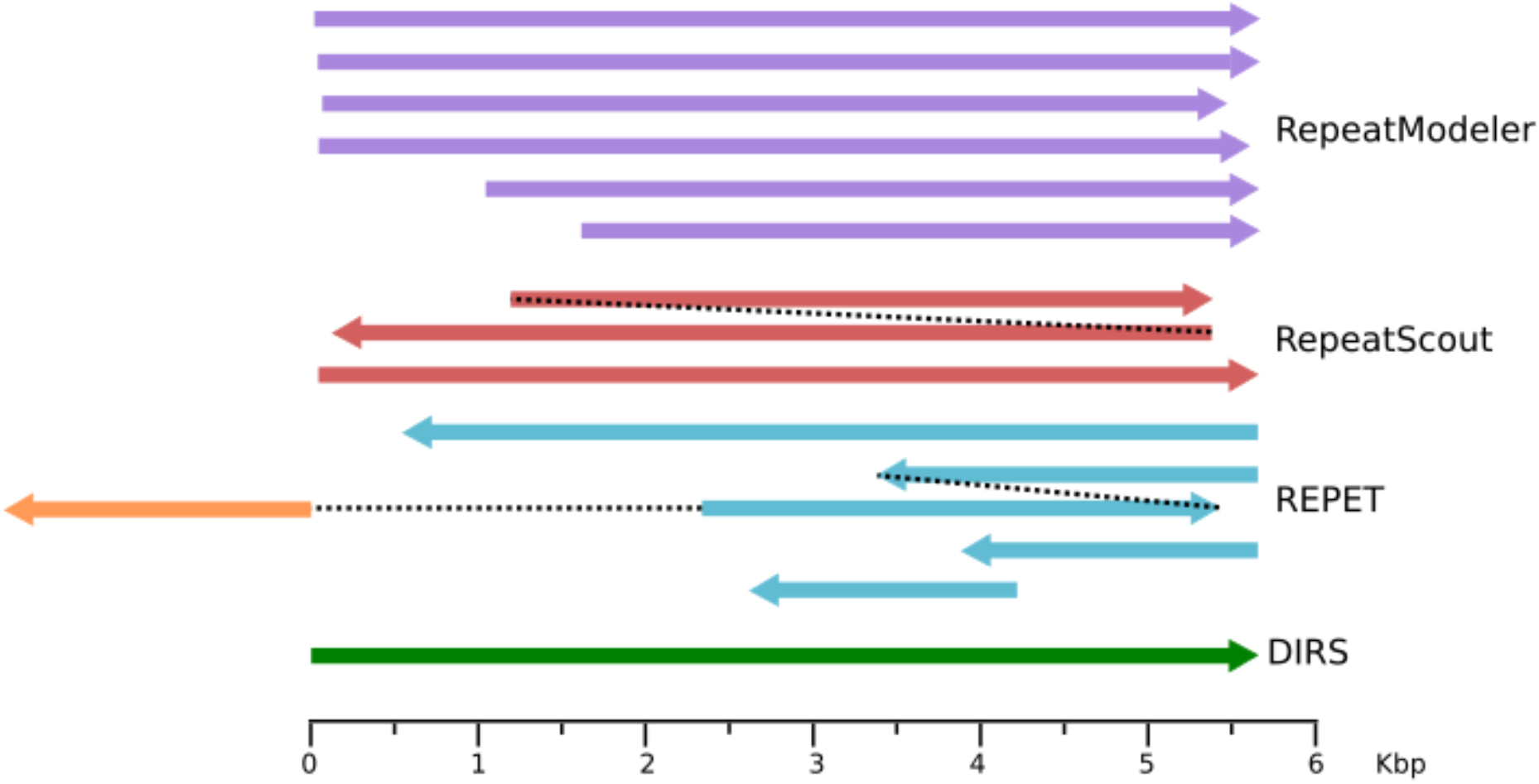
Consensus sequences generated by each software for a single TE from the DIRS family. The length and position reflect the mapping to the TE, black dotted lines show the continuation of the same model, the orange segment represents a fragment coming from another TE (TRANSIB).

Another model is a hybrid TE consisting of two overlapping fragments of DIRS element in a head-to-tail orientation with a fragment of TRANSIB transposon. The fourth model closely resembles the original TE but is truncated by about 240 nucleotides at its 5’ end (see Figure 5).

In Figure 4 we present models generated for another TE, namely Polinton-1_DR (Kapitonov and Jurka 2006). The full-length transposon is 18.5 kb, including 350 bp terminal inverted repeats. All three software compared in our study reported few models for this TE but none of the models recovered the full length TE (see Figure 4 and Table S1). Interestingly, both REPET and RepeatModeler generated similar close-to-full-length models that are missing one of the inverted repeats, while RepeatScout’s longest model misses both inverted repeats. However, inverted repeats were reported as separate models by each of the program.

In the real data from model organisms, RepeatScout created the highest number of models with almost three-thousand consensuses for zebrafish chromosome 1 (see Table 3). This is in contrast to the simulated data where RepeatScout generated the least number of models. REPET lies on the other extreme of the spectrum with just 65 TE models for the human chromosome 21, including Alu model that was missing from the simulated data analysis.

**Table 3.** Number of models generated by three software in real sequence analysis and number of TE-families annotated in publicly available data on the same sequences.

| Software | Zebrafish chromosome 1 | Fruit fly genome | Human chromosome 21 |
| --- | --- | --- | --- |
| RepeatScout | 2919 | 2593 | 464 |
| RepeatModeler | 1779 | 686 | 428 |
| REPET | 342 | 557 | 65 |
| Public annotation | 875 | 219 | 979 |

Interestingly, this chromosome is annotated with almost 1000 different TE families. The smaller number of models generated for human data might be linked to the smaller sequence data compared to the two other datasets. However, based on the TE-annotation, the total length of TEs in the fruit fly genome is comparable to the total length of TEs in human chromosome 21, 24 MB versus 20 Mb, respectively. In general, the real data seem to harbor more versatile repertoire of TEs than our simulated data resulting in many more TE models (compare Table 2 and Table 3).

To investigate this matter further, we compared TE models created by the software analyzed with TE families annotated in the sequences used for the benchmarking. We simply run RepeatMasker with the created TE libraries as a query. Interestingly, in all cases there were many consensus sequences (TE models) that didn’t produce any significant hits in the RepeatMasker analysis, suggesting that there might still be undiscovered repeats in the analyzed data (see Table 4).

**Table 4.**
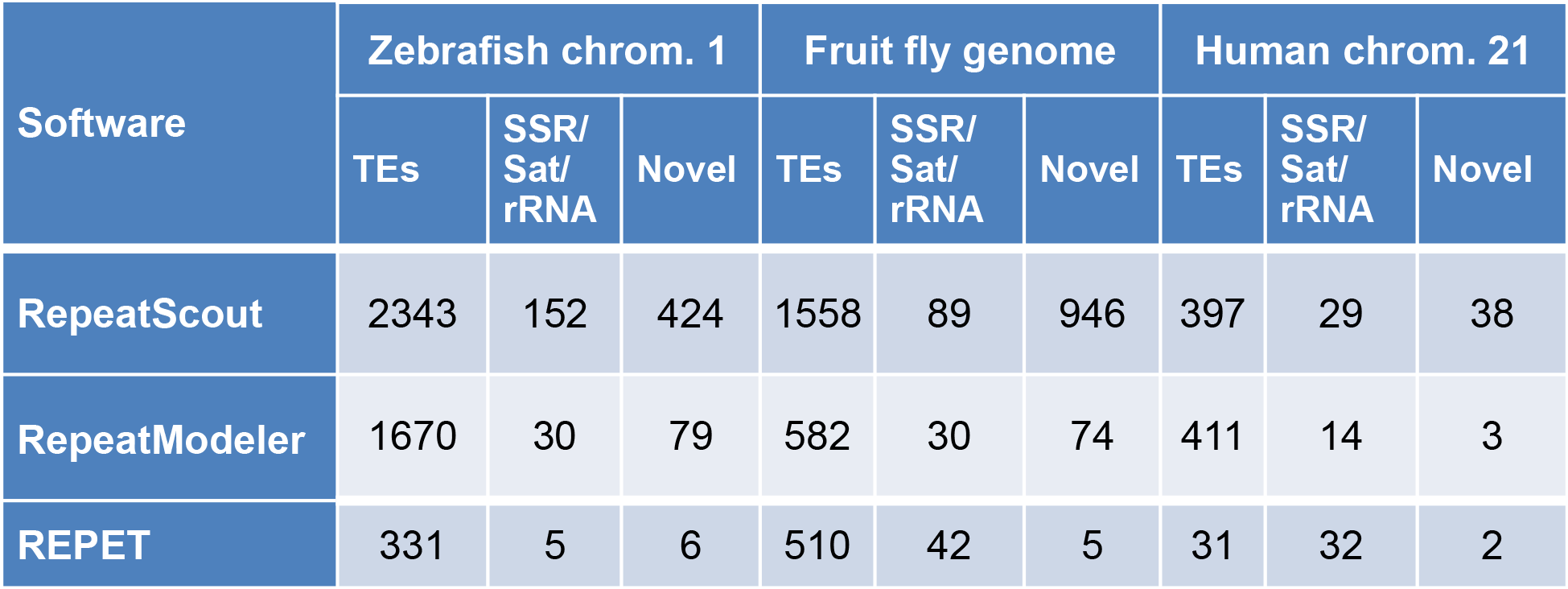
General annotation of models built by analyzed software using RepeatMasker and cognate libraries. In the “Novel” column we list the number of models that didn’t produce any individual matches during RepeatMasker run.

| Software | Zebrafish chrom. 1 |  |  | Fruit fly genome |  |  | Human chrom. 21 |  |  |
| --- | --- | --- | --- | --- | --- | --- | --- | --- | --- |
|  | TEs | SSR/<br>Sat/<br>rRNA | Novel | TEs | SSR/<br>Sat/<br>rRNA | Novel | TEs | SSR/<br>Sat/<br>rRNA | Novel |
| RepeatScout | 2343 | 152 | 424 | 1558 | 89 | 946 | 397 | 29 | 38 |
| RepeatModeler | 1670 | 30 | 79 | 582 | 30 | 74 | 411 | 14 | 3 |
| REPET | 331 | 5 | 6 | 510 | 42 | 5 | 31 | 32 | 2 |

### Individual repeats annotation

To get a better idea of the different results of the *de novo* annotation obtained by the six tools used, we plotted the coordinates of each one in tracks along with the reference annotation, as shown in Figure 6. Simply by visualizing the results it is quite evident that there is a tendency to get a fragmented annotation when using k-mer counting tools, particularly Pclouds and phRaider. Red uses a smoothing function to merge nearby high frequency k-mers, giving less fragmented results, as shown in Figure 6 and 7. As it was expected due to the methodology used, the best results for detecting transposons were obtained by software that calculate TE-models, but as it is shown more in detail in Figure 7, most of the predictions are fragmented TEs, annotations without clear borders, or missing some smaller or incomplete elements.

**Figure 6.**
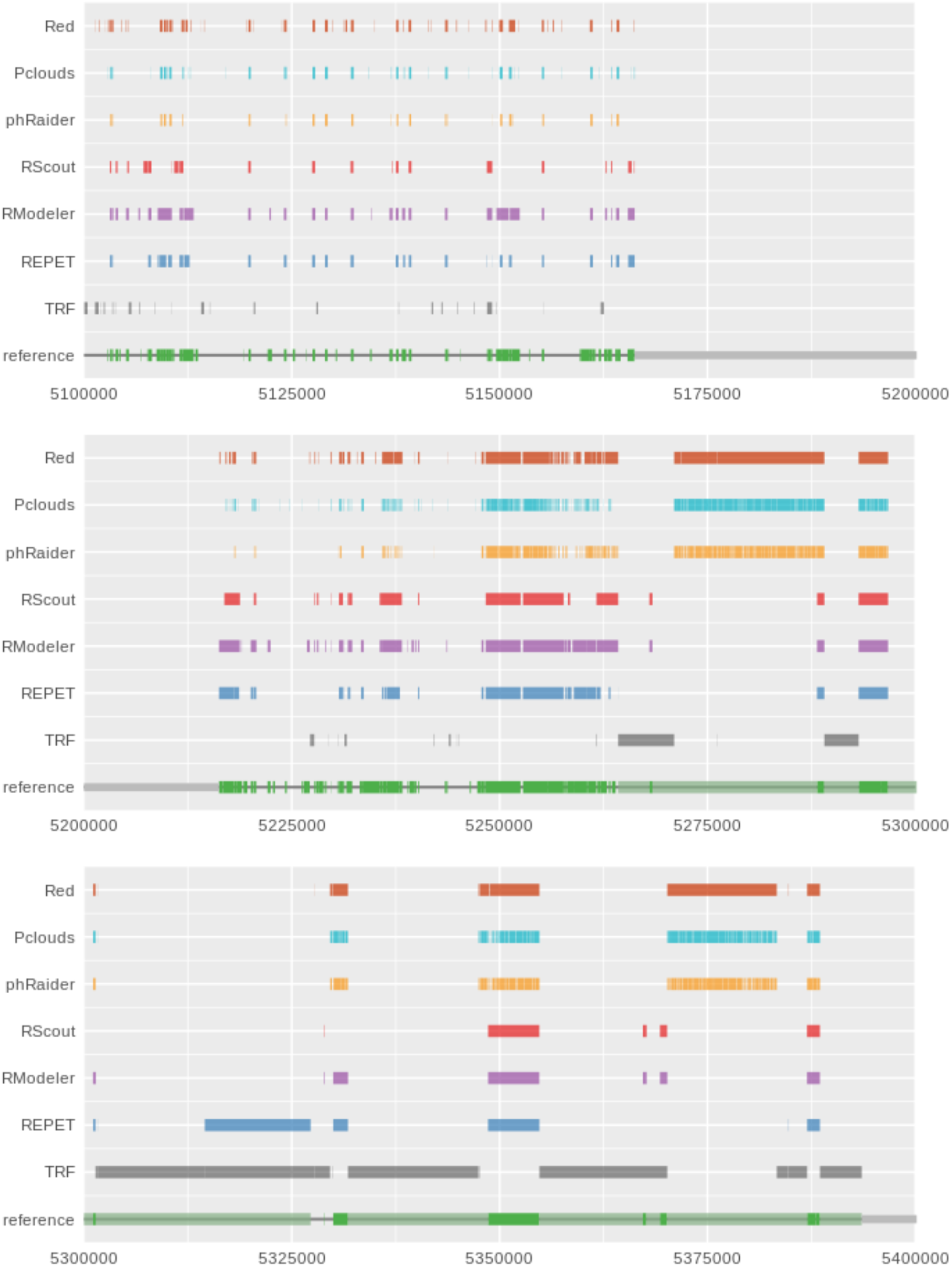
Different tracks of the coordinates obtained from a *de novo* identification of transposons using six different software tools for detecting interspersed repeats. In the reference track, green blocks are transposons.

**Figure 7.**
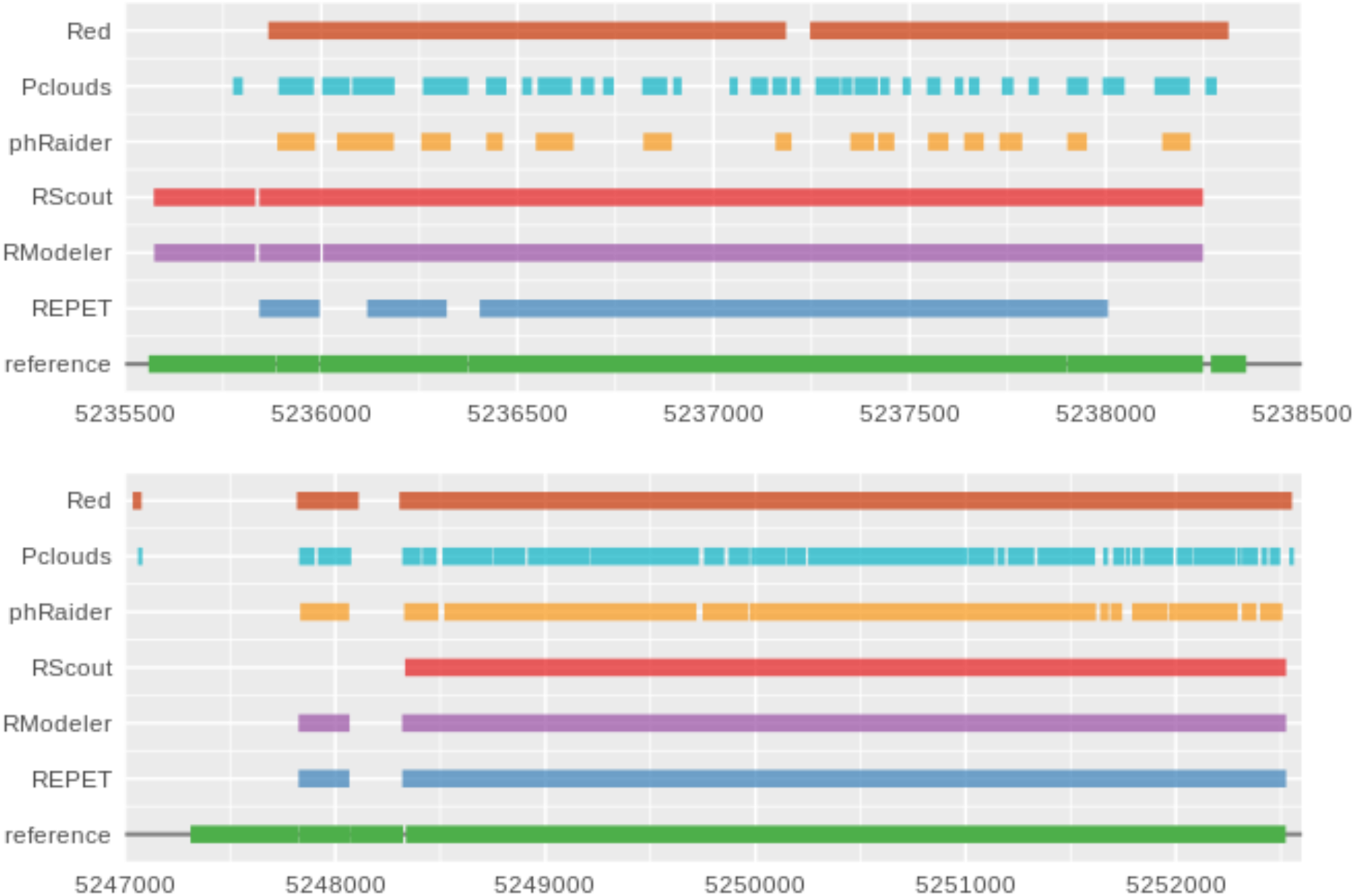
Comparison of the fragmentation of results in a region of the human chromosome 21. In each track are the predictions obtained from each tool and in green the reference. Notice how most of the results are usually incomplete or fragmented.

For model-building software, in most cases RepeatModeler got the best results, but RepeatScout obtained also comparable results. REPET failed in some scenarios and particularly with short divergent fragments (Alu elements). However, it must be noticed that it failed only in one of the twenty cases and performed quite well on a real data. One of the reasons could be that this software was developed with the idea to be used with large genomes and here all the tests were run with sequences of around 100 MB. We compared the annotation results obtained using simulated sequences with known identities between TEs ranging from 60% to 100% and then compared the coverage of the annotation in relation to it, as is shown in Figure 6. It is expected that TEs with higher identities are detected more precisely. Indeed TEs with a higher identity are better detected by all software and the differences seen are inherent to the performance of each tool. For k-mer counting software, Red performs significantly better than the rest, e.g at 70% identity, Red detects approximately 85% of the TE regions. Meanwhile, Pclouds detects about 60% and phRaider 25% (Figure 8). The model-building software display a much better and uniform performance and are less affected by more divergent TEs (Figure 8). When we also consider the different TE orders annotated and the proportion of coverage for the TEs of each order, we observe that there’s no significant difference between them and all the software have a consistent performance (supplementary material, Figure S1).

**Figure 8.**
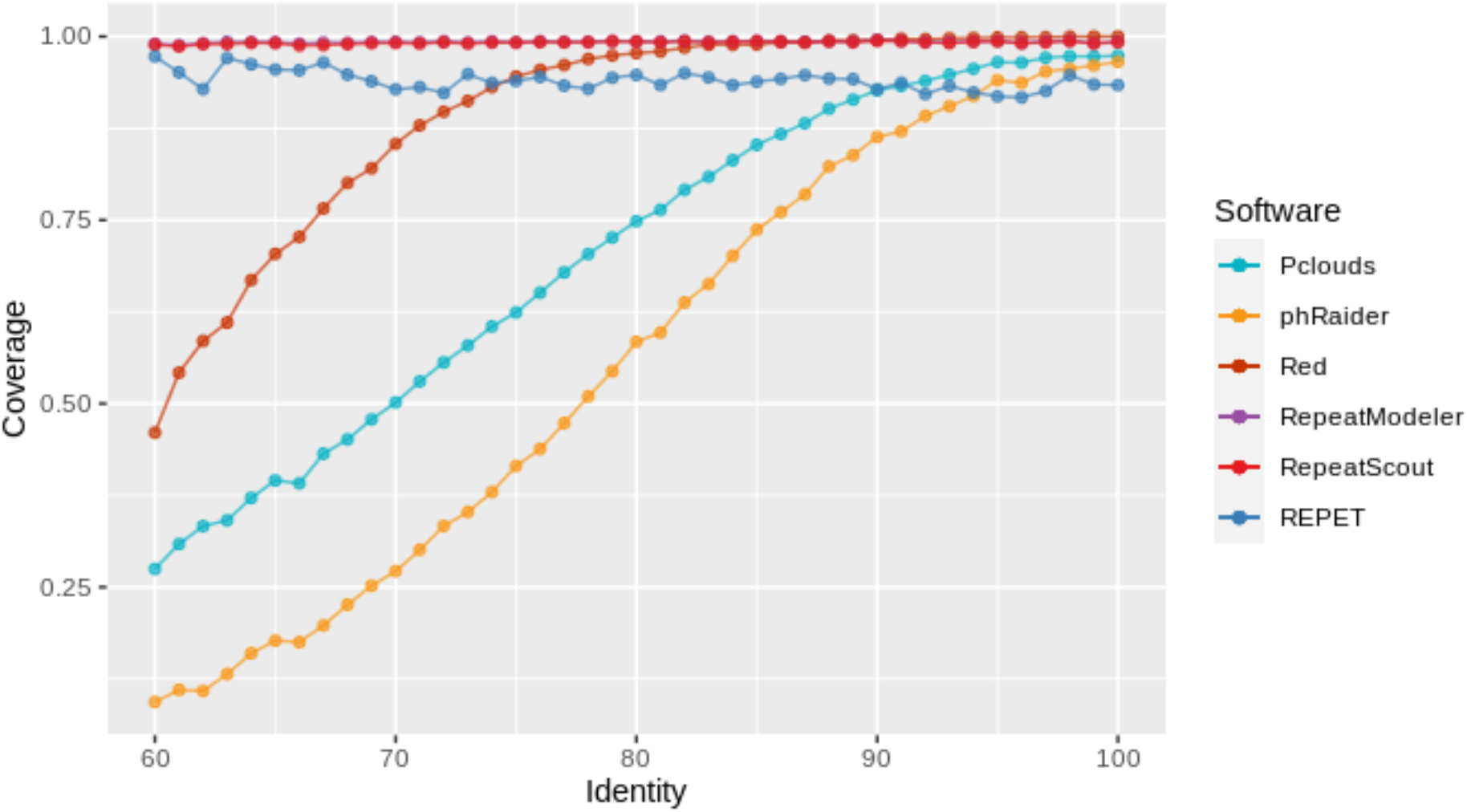
Coverage of TEs in simulated sequence in relation to their average identity. In k-mer counting software there’s a great drop in the detection of more divergent TEs, this behavior is not seen when using model-builders. RepeatModeler and RepeatScout results are almost identical and they are overlapped in this plot.

We also analyzed how well computed libraries can detect individual TEs and masked genomic sequences. To do that, we run RepeatMasker with genomic sequences as queries and build by different software libraries as references. In most cases RepeatScout libraries gave the results closest to the current expert annotation of cognate genomic sequence (see Table 5).

**Table 5.** Number of nucleotides detected as TEs by software analyzed. Numbers in parentheses represent fraction of the sequence covered by those TEs.

| Reference library | Zebrafish chromosome 1 | Fruit fly genome | Human chromosome 21 |
| --- | --- | --- | --- |
| RepeatScout | 26,447,536<br>(44.39%) | 23,540,405<br>(17.11%) | 12,266,284<br>(26.26%) |
| RepeatModeler | 16,561,994<br>(27.80%) | 15,919,335<br>(11.57%) | 13,551,826<br>(29.01%) |
| REPET | 18,712,280<br>(31.41%) | 22,412,191<br>(16.29%) | 8,560,157<br>(18.33%) |
| Original annotation | 26,465,290<br>(44.42%) | 23,990,534<br>(17.44%) | 17,188,977<br>(36.80%) |
| Size of analyzed sequence | 59,578,282 | 137,547,960 | 46,709,983 |

Interestingly, none of the libraries seem to work well with the human chromosome 21 data masking only between 50 and 79 per cent of the originally masked sequences. However, if we look at the number of annotated TEs, the situation is not looking as bad (see Table 6). This is probably due the fact that all the software used here produced libraries with rather fragmented models as compared to curated data (see discussion above).

**Table 6.** Number of individual TEs annotated by RepeatMasker using different reference libraries.

| Reference library | Zebrafish chromosome 1 | Fruit fly genome | Human chromosome 21 |
| --- | --- | --- | --- |
| RepeatScout | 143463 | 55799 | 46628 |
| RepeatModeler | 86813 | 30754 | 49621 |
| REPET | 71891 | 38154 | 30838 |
| Original annotation | 107997 | 38282 | 53540 |

Finally we evaluated the performance of each tool against the datasets using the Matthews Correlation Coefficient (Figure 9). To evaluate the results it is important to consider not only the raw performance of each tool but also the difficulty to run, configurability, and speed. K-mer counting software usually only accept a few parameters such as k-mer length, minimum frequency, and length; but these tools are usually very easy to run and require little computing power while being incredibly fast. However, one of the performance downsides can be the requirement to store large data structures in memory. In this category, with defaults parameters Red outperformed pClouds and phRaider and this can be explained by the fact that Red merges nearby k-mers more frequently than the others, giving less fragmented results. Model-building software employ a strategy that requires much more computational resources. They are also more time consuming and can be more complex to install, configure, and run. In our tests RepeatModeler obtained the best results, but it is interesting to notice that RepeatScout that is part of RepeatModeler pipeline is faster than the latter but obtained almost as good results as the pipeline. However, it should pointed out that RepeatModeler tries to classify calculated models into repeat families, something that RepeatScout does not. Hence, execution time difference is rather expected than surprising. REPET on the other hand was probably not the best tool for this type of analysis. REPET has a default configuration but also different tools can be added to the pipeline and each step is highly configurable, but one of the downsides is that it can be very complex to configure and run for an unexperienced user.

**Figure 9.**
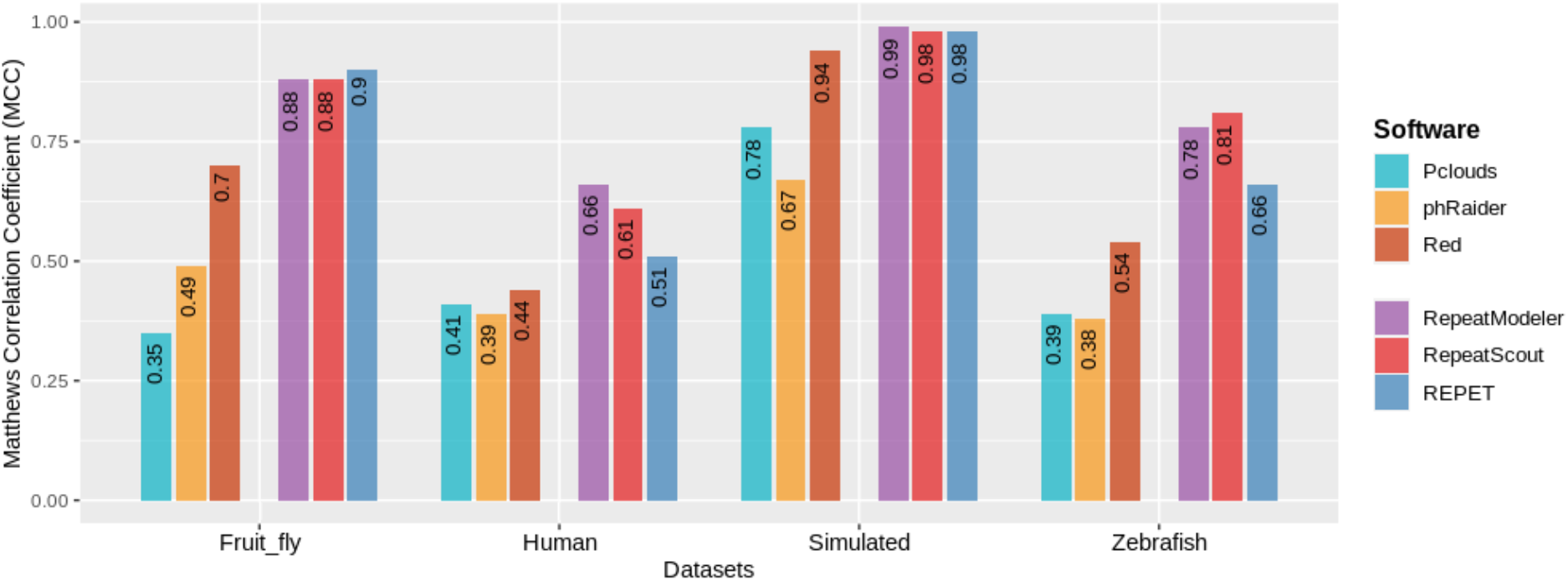
Matthews Correlation Coefficient values showing the performance of each tool tested with the datasets tested.

## Discussion

Presented here benchmark is not very optimistic but how does it measure against other similar studies. Unfortunately, the comparison is not very easy. First of all, there are not many independent benchmarking tests perform in past (Saha et al. 2008; Ou et al. 2019). Although, usually some benchmarking are presented with the original publication of a software, they cannot be completely trusted as they might be tuned to a specific software. Moreover, each study includes different set of the software and we didn’t find any single paper that discussed the same six programs that we benchmarked here. Ou et al. employed five programs of which three (RepeatScout, RepeatModeler, and Red) were benchmarked by us as well. However, in this paper benchmarking was used to compare these software with EDTA, a pipeline created by the authors and as such cannot be treated as an independent analysis. We found only one, truly independent benchmarking study but it was published over a decade ago and only RepeatScout is a mutual software with our study (Ou et al. 2019). Although RepeatModeler is most frequently compared software, surprisingly it is missing from Ou et al. analysis. Another difficulty is that each study used different data sets to evaluate *de novo* TE-detection software and surprisingly simulated data usually was not included. Similarly to our approach the datasets varied in size but none of the tools was tested on a sequence of gigabases scale. Nevertheless, below we try to compare our findings with those previously published.

First of all, it is clear that in most studies RepeatModeler gives the best results. In the original paper Flynn et al. divided computed models into perfect, good, present, and not found (Flynn et al. 2020). Although this classification might be a bit misleading, e.g. “perfect” models might only 95% identical with the reference consensus, we looked at our results the same way (see Fig. 10). Interestingly, although in both studies *D. melanogaster* genome was subjected to the analysis, the results were not the same. Most likely, it is due to the fact that we used default parameters of RepeatModeler.

**Figure 10.**
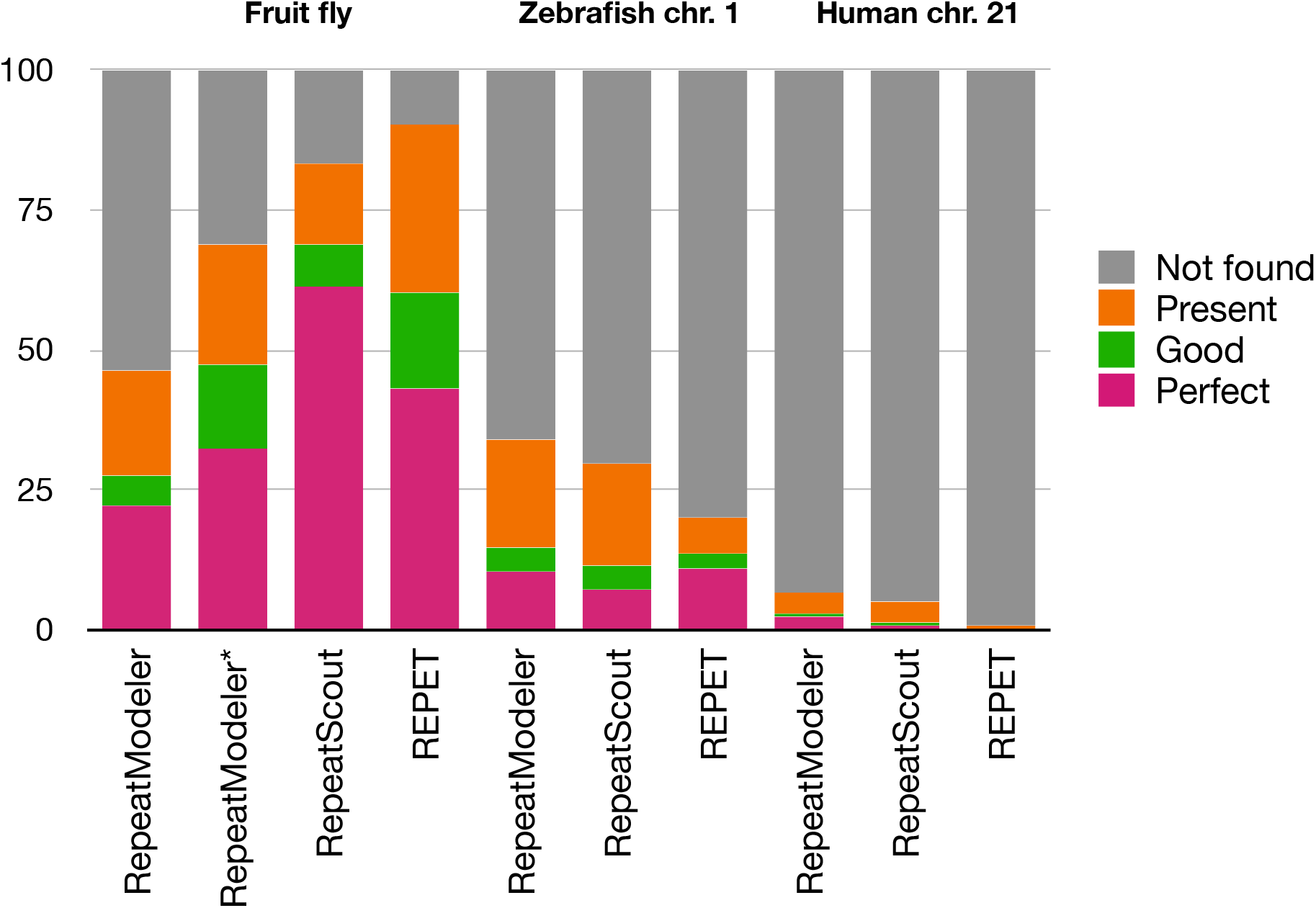
Summary of models’ accuracy defined by different software and data sets. RepeatModeler* data taken from Flynn et al. 2020. Please refer to Fig. 3 in this paper for categories definition.

As mentioned above Saha et al. benchmarked six programs but only RepeatScout was mutual software between their and our study (Saha et al. 2008). Moreover, they run tests only on different rice genome data varying from 3 to 27 Mb only with published annotation assumed as a base line (golden standard) and only sensitivity was reported. For unknown reason, different programs were run on different datasets. Nevertheless, RepeatScout performed the worst with 3 Mb sequence data with only 26.2% sensitivity but performed much better when run on a larger 27.8 Mb dataset with sensitivity increased to 84.3%. The other software used on both datasets (RepeatFinder) behaved similarly, 32.7% and 85.3% sensitivity, respectively.

These results are similar to our observations that *de novo* repeat finding programs perform better when a larger dataset is used to build a TE library (see Table 5).

Based on our simulated data analysis it is clear that none of the analyzed software is able to compute a repeat consensus sequence perfectly. While the sequence of a repeat can be recovered with confidence, the structure of the repeat should be inspected manually and edited accordingly. This is especially important for longer transposons. In general, for a fast assessment of interspersed repeats, Red can be useful, acknowledging of course its limitations when it comes to low complexity sequences. For more in depth studies with small genomes, RepeatModeler seems to be the best option. It is also interesting to note that RepeatScout has a really good performance if we take into consideration speed and computational requirements. However, the situation may not be as bad as it sounds. For some tasks such as repeat masking this might be satisfactory. Nevertheless, to fully understand biology and evolution of transposable elements in a given genome automatic approach is not sufficient but it should be a good starting point.

## Conclusions

We tested a number of tools for *de novo* detection of TEs. The results were compared using the MCC against a reference of annotated TEs. As expected, model-builders performed better than k-mer counting software, with RepeatModeler beating competitors in most datasets. However, even for RepeatModeler, the results are far from satisfactory based on the reference annotation. There is a tendency for most tools to identify TE-regions in a fragmented manner and it is also frequent that small TEs or fragmented TEs are not detected. We recognize that some of the results obtained may be improved by fine tuning of parameters; some tools like REPET are fully customizable and more tools can be added to the pipeline, although this can be challenging for most users. In conclusion, the contemporary tools for *de novo* detection of TEs benchmarked here are far from being perfect and the identification of TEs is still a challenging endeavor as it requires a significant manual curation by an experienced expert. It seems that for *de novo* detection of TEs extensive manual curation and using multiple tools for confirmation of the results obtained is necessary. We also found that MCC can be used as a fast and reliable test to compare the performance of these software and can give a general idea of which tool is best suited for each task.

## Declarations

### Ethics approval and consent to participate

Not applicable.

### Consent for publication

Not applicable.

### Availability of data and materials

Scripts and simulated sequences used for the software evaluation are deposited at https://github.com/IOB-Muenster/denovoTE-eval.

### Funding

This work was partially funded by the DAAD Research Grants - Doctoral Programmes in Germany, 2018/19 (57381412) to MR and internal funds of the Institute of Bioinformatics.

### Competing interests

The authors declare that they have no competing interests.

### Authors’ contributions

WM initiated and designed the project. MR executed the project and wrote a draft manuscript. Both authors worked on the final manuscript.

## Acknowledgements

Not applicable.

## Supplementary materials

**Table S1.** TEs inserted into simulated data. Transposon ids after Dfam.

| TE_id | Count | Average identity | Standard deviation | Number of indels | Target site duplication | Length | Number of truncated elements | Number of nested element |
| --- | --- | --- | --- | --- | --- | --- | --- | --- |
| **AluY** | 730 | 87 | 12 | 16 | y | 311 | 67 | 11 |
| **BEL12-I_DR** | 550 | 89 | 12 | 10 | y | 7730 | 80 | 12 |
| **Copia1_DM** | 680 | 87 | 14 | 10 | y | 4530 | 65 | 10 |
| **DIRS-1_DR** | 710 | 85 | 15 | 12 | n | 5634 | 70 | 5 |
| **G5_DM-Jockey** | 590 | 91 | 10 | 15 | y | 4856 | 75 | 16 |
| **GYPSY8_DM** | 470 | 84 | 10 | 15 | y | 4955 | 74 | 12 |
| **hAT-9_DR** | 480 | 82 | 12 | 11 | y | 2386 | 65 | 13 |
| **Helitron-1N3_DR** | 550 | 83 | 10 | 11 | n | 1843 | 65 | 10 |
| **HERVL** | 590 | 88 | 14 | 14 | y | 6542 | 50 | 7 |
| **L1-4_DR** | 620 | 78 | 9 | 16 | y | 5548 | 72 | 13 |
| **Merlin1_HS** | 580 | 80 | 11 | 13 | y | 1175 | 60 | 18 |
| **Ngaro1_DR** | 560 | 86 | 12 | 11 | n | 6578 | 73 | 9 |
| **Penelope-1_DR** | 810 | 85 | 11 | 20 | y | 4446 | 60 | 11 |
| **piggyBac-N1_DR** | 610 | 80 | 10 | 19 | y | 1005 | 64 | 20 |
| **Polinton-1_DR** | 420 | 81 | 11 | 20 | n | 18485 | 70 | 10 |
| **RTE1** | 570 | 90 | 13 | 18 | y | 3291 | 71 | 11 |
| **SINE3-1** | 580 | 85 | 13 | 10 | y | 590 | 70 | 7 |
| **TC1-2_DM** | 590 | 83 | 14 | 12 | y | 1644 | 70 | 16 |
| **TRANSIB1** | 670 | 81 | 10 | 12 | y | 3014 | 68 | 9 |
| **Turmoil2** | 650 | 79 | 9 | 10 | y | 6999 | 69 | 15 |

**Table S2.** Comparison of the consensus models obtained for each single TE inserted in the simulated sequence. With * are indicated models were the models include a duplication or an extension of the same TE and with # are indicated models that are the combination of two TEs which originated from nested TEs.

| Original TE-families |  | Length of the models created |  |  |
| --- | --- | --- | --- | --- |
| TE | Length | RepeatScout | RepeatModeler | REPET |
| AluY | 311 | 308 | 345<br>320<br>310 | - |
| BEL12 | 7730 | 7724 | 7751<br>7730<br>6413 | 11148*<br>7750<br>2769 |
| Copia | 4530 | 8843* | 4532<br>4514<br>4326<br>4076 | 4652<br>4314 |
| DIRS | 5654 | 5644<br>9385* | 5672<br>5653<br>5610<br>5445<br>4633<br>4062 | 7134#<br>5401<br>1824<br>1727 |
| G5 | 4856 | 4854 | 4851<br>4843<br>4836<br>3151 | 7502 *<br>5232<br>5197<br>4904<br>3967<br>2359 |
| GYPSY | 4955 | 4360<br>3920 | 4938<br>4012<br>3951<br>3373 | 5068<br>4682<br>2061<br>1445<br>1256 |
| hAT | 2386 | 2380 | 2380<br>2380 | 2426 |
| Helitron | 1843 | 1839<br>1472 | 1832<br>1823<br>1234 | 1907<br>1906<br>1744<br>871<br>633 |
| TE | Length | RepeatScout | RepeatModeler | REPET |
| HERVL | 6542 | 12634* | 6549<br>6523<br>6094 | 11495#<br>6553<br>3886<br>3169 |
| L1 | 5548 | 4460<br>3379<br>2415 | 5496<br>5488<br>3386<br>2588<br>2478<br>1289 | 5784<br>5605<br>5443<br>2855<br>2142<br>1794<br>1639<br>1179<br>667 |
| Merlin | 1175 | 1168 | 1172<br>1167<br>744 | 1231<br>1216<br>1089<br>629 |
| Ngaro | 6578 | 6571<br>6020 | 6568<br>6566<br>6563<br>6156 | 7798#<br>6652<br>6379<br>2487<br>2445 |
| Penelope | 4446 | 4437<br>2276 | 4432<br>4407<br>4189<br>3910<br>3198<br>3908 | 5077<br>3251<br>812 |
| piggyBac | 1005 | 997 | 1002<br>990<br>840 | 1067<br>1055<br>966<br>641<br>477 |
| Polinton | 18485 | 14288<br>10127<br>4536<br>1513 | 17626<br>13482<br>11066<br>9356<br>8330<br>6655<br>372<br>169 | 16912<br>16015<br>15368<br>10418<br>6631<br>2793<br>1690<br>1216 |
| TE | Length | RepeatScout | RepeatModeler | REPET |
| RTE1 | 3291 | 3287 | 3286<br>3270<br>3212 | 3343<br>3200<br>3128<br>1222 |
| SINE3 | 590 | 587 | 587<br>581<br>578 | 624<br>559 |
| TC1 | 1644 | 1634 | 1636<br>1627<br>1472 | 1689<br>771 |
| TRANSIB | 3014 | 3002 | 2995<br>2546<br>2378 | 3094<br>2858<br>1166 |
| Turmoil | 6999 | 6676 | 7016<br>6985<br>6974<br>6970<br>4201 | 7113<br>6598<br>5956<br>5834#<br>1757<br>1381<br>1100 |

**Figure S1.**
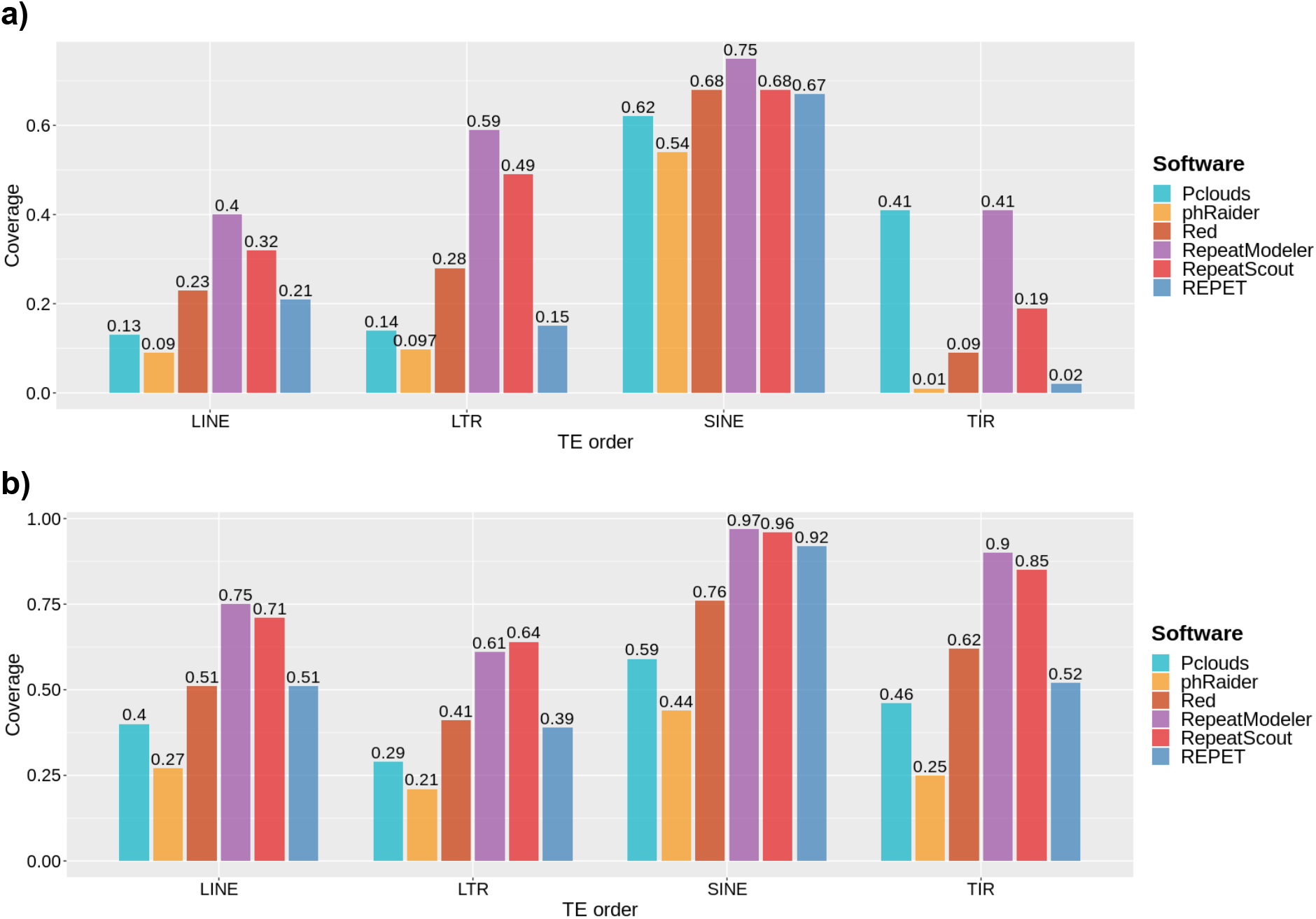
Coverage of each TE order in the human (a) and zebrafish (b) dataset.

## Notes

https://github.com/IOB-Muenster/denovoTE-eval

### Competing Interest Statement

The authors have declared no competing interest.

## Bibliography

Bao, W.D., K.K. Kojima, and O. Kohany, 2015 Repbase Update, a database of repetitive elements in eukaryotic genomes. Mobile DNA 6.

Benson, G., 1999 Tandem repeats finder: a program to analyze DNA sequences. Nucleic Acids Research 27 (2):573–580.

Biemont, C., 2010 A Brief History of the Status of Transposable Elements: From Junk DNA to Major Players in Evolution. Genetics 186 (4):1085–1093.

Boughorbel, S., F. Jarray, and M. El-Anbari, 2017 Optimal classifier for imbalanced data using Matthews Correlation Coefficient metric. Plos One 12 (6).

de Koning, A.P.J., W.J. Gu, T.A. Castoe, M.A. Batzer, and D.D. Pollock, 2011 Repetitive Elements May Comprise Over Two-Thirds of the Human Genome. Plos Genetics 7 (12).

Flutre, T., E. Duprat, C. Feuillet, and H. Quesneville, 2011 Considering Transposable Element Diversification in De Novo Annotation Approaches. Plos One 6 (1).

Flynn, J.M., R. Hubley, C. Goubert, J. Rosen, A.G. Clark et al., 2020 RepeatModeler2 for automated genomic discovery of transposable element families. Proceedings of the National Academy of Sciences of the United States of America 117 (17):9451–9457.

Gao, C.H., M.L. Xiao, X.D. Ren, A. Hayward, J.M. Yin et al., 2012 Characterization and functional annotation of nested transposable elements in eukaryotic genomes. Genomics 100 (4):222–230.

Girgis, H.Z., 2015 Red: an intelligent, rapid, accurate tool for detecting repeats de-novo on the genomic scale. Bmc Bioinformatics 16.

Gu, W.J., T.A. Castoe, D.J. Hedges, M.A. Batzer, and D.D. Pollock, 2008 Identification of repeat structure in large genomes using repeat probability clouds. Analytical Biochemistry 380 (1):77–83.

Haeussler, M., A.S. Zweig, C. Tyner, M.L. Speir, K.R. Rosenbloom et al., 2019 The UCSC Genome Browser database: 2019 update. Nucleic Acids Research 47 (D1):D853–D858.

Hoen, D.R., G. Hickey, G. Bourque, J. Casacuberta, R. Cordaux et al., 2015 A call for benchmarking transposable element annotation methods. Mobile DNA 6.

Hubley, R., R.D. Finn, J. Clements, S.R. Eddy, T.A. Jones et al., 2016 The Dfam database of repetitive DNA families. Nucleic Acids Research 44 (D1):D81–D89.

Jurka, J., V.V. Kapitonov, O. Kohany, and M.V. Jurka, 2007 Repetitive sequences in complex genomes: Structure and evolution. Annual Review of Genomics and Human Genetics 8:241–259.

Kapitonov, V.V., and J. Jurka, 2006 Self-synthesizing DNA transposons in eukaryotes. Proc Natl Acad Sci U S A 103 (12):4540–4545.

Kubiak, M.R., and I. Makalowska, 2017 Protein-Coding Genes’ Retrocopies and Their Functions. Viruses 9 (4).

Makalowski, W., 2000 Genomic scrap yard: how genomes utilize all that junk. Gene 259 (1-2):61–67.

Makalowski, W., V. Gotea, A. Pande, and I. Makalowska, 2019 Transposable Elements: Classification, Identification, and Their Use As a Tool For Comparative Genomics. Methods Mol Biol 1910:177–207.

Ohno, S., 1973 So much “junk” DNA in our genome, pp. 366–370 in Evolution of Genetic Systems: Brookhaven Symposia in Biology., edited by H. Smith. Gordon and Breach, New York.

Ou, S., W. Su, Y. Liao, K. Chougule, J.R.A. Agda et al., 2019 Benchmarking transposable element annotation methods for creation of a streamlined, comprehensive pipeline. Genome Biol 20 (1):275.

Price, A.L., N.C. Jones, and P.A. Pevzner, 2005 De novo identification of repeat families in large genomes. Bioinformatics 21:I351–I358.

Quesneville, H., C.M. Bergman, O. Andrieu, D. Autard, D. Nouaud et al., 2005 Combined evidence annotation of transposable elements in genome sequences. Plos Computational Biology 1 (2):166–175.

Ricker, N., H. Qian, and R.R. Fulthorpe, 2012 The limitations of draft assemblies for understanding prokaryotic adaptation and evolution. Genomics 100 (3):167–175.

Saha, S., S. Bridges, Z.V. Magbanua, and D.G. Peterson, 2008 Empirical comparison of ab initio repeat finding programs. Nucleic Acids Research 36 (7):2284–2294.

Schaeffer, C.E., N.D. Figueroa, X.L. Liu, and J.E. Karro, 2016 phRAIDER: Pattern-Hunter based Rapid Ab Initio Detection of Elementary Repeats. Bioinformatics 32 (12):209–215.

Schnable, P.S., D. Ware, R.S. Fulton, J.C. Stein, F. Wei et al., 2009 The B73 maize genome: complexity, diversity, and dynamics. Science 326 (5956):1112–1115.

Smit, A., R. Hubley, and P. Green, 2013-2015 RepeatMasker Open-4.0.

Wicker, T., F. Sabot, A. Hua-Van, J.L. Bennetzen, P. Capy et al., 2007 A unified classification system for eukaryotic transposable elements. Nature Reviews Genetics 8 (12):973–982.

